# Modeling the control of bacterial infections via antibiotic-induced proviruses

**DOI:** 10.1101/706796

**Authors:** Sara M. Clifton, Ted Kim, Jayadevi H. Chandrashekhar, George A. O’Toole, Zoi Rapti, Rachel J. Whitaker

## Abstract

Most bacteria and archaea are infected by latent viruses that change their physiology and responses to environmental stress. We use a population model of the bacteria-phage relationship to examine the role that latent phage play on the bacterial population over time in response to antibiotic treatment. We demonstrate that the stress induced by antibiotic administration, even if bacteria are resistant to killing by antibiotics, is sufficient to control the infection under certain conditions. This work expands the breadth of understanding of phage-antibiotic synergy to include both temperate and chronic viruses persisting in their latent form in bacterial populations.

**Importance:** Antibiotic-resistance is a growing concern for management of common bacterial infections. Here we show that antibiotics can be effective at sub-inhibitory levels when bacteria carry latent phage. Our findings suggest that specific treatment strategies based on the identification of latent viruses in individual bacterial strains may be an effective personalized medicine approach to antibiotic stewardship.

## INTRODUCTION

A world-wide growth of antibiotic resistance threatens the efficacy of antibiotic treatments for common infections, driving medical professionals to seek alternative treatments [1]. Infections by *Pseudomonas aeruginosa* alone represent about 10% of nosocomial infections, are a leading cause of death among patients with cystic fibrosis (CF), and have been deemed a serious threat on the United States Centers for Disease Control watch list for antibiotic resistance [2–4]. Despite the increasing trend of multidrug resistance, antibiotic regimes remain the consensus first treatment for *P. aeruginosa* infection [5]. As a last resort and as an attempt to prevent the evolution of resistance in *P. aeruginosa*, clinicians have turned to combination therapies [6] with bacteriophage (viruses) and antibiotics to treat recalcitrant bacteria.

Synergy between phage and antibiotic treatment (PAS) is now rising in interest for treatment of *P. aeruginosa* and other recalcitrant bacteria [7–9]. Combination phage therapy uses viruses that kill bacteria (often in phage cocktails) and different types of antibiotics either at the same time or in series to clear bacteria and prevent the evolution of new resistant phenotypes [10–18]. Although preexisting proviruses are highly prevalent in *P. aeruginosa* infections and appear to be induced by certain antibiotic treatments, synergy has not been considered in the context of temperate virus induction. Here, we investigate the role phages play during antibiotic treatment when they are already present in the system. We show that, even without deliberate phage therapy, phages may play a critical role in antibiotic treatment especially if the bacteria are antibiotic resistant.

## BACKGROUND

Bacteriophages are viruses that infect bacteria and hijack cell functions in order to reproduce. Just as bacteria have evolved many strategies to evade infection, phages have developed multiple strategies to circumvent cell defenses. Phages can be characterized by their lifestyles (obligately lytic, temperate, or chronic) within the host [19]. Lytic viruses replicate within the host and kill host cells by bursting them open to release new particles. Temperate viruses have a lytic cycle but can also integrate into host genomes where they remain latent until they are induced to replicate [19]. In chronic infection, productive host cells shed new phages that bud from the cell without killing the bacterium [20]. Both temperate and chronic viruses have a lysogenic (latent lytic or latent chronic) cycle in which phage DNA is incorporated into the bacterium’s genome, and the cell transmits the phage’s genetic material (prophage) to daughter cells vertically [21].

Comparative genomics among closely related bacterial strains has uncovered a plethora of proviruses of both temperate and chronic lifestyles [22–24]. The large genome of the opportunistic pathogen *P. aeruginosa* is no exception [25–27]. Each sequenced strain reveals multiple proviral genomes of both the temperate and chronic lifestyles, each in both active and inactive (latent) forms [28]. These proviruses change bacterial fitness and environmental response, sometimes conferring competitive advantage, virulence, and antibiotic resistance [29–32].

Stressful environmental conditions (e.g., radiation, heat, sublethal antibiotics) may trigger the cell to induce latent prophage and begin phage production [33–37]. The induction of such latent phages is proposed to be one of the mechanisms behind the synergistic effect of antibiotics and phage infection [37, 38]. The environmental conditions, especially dynamic antibiotic dosing regimes, under which these phage types may coexist are not well understood. We therefore develop a population model to understand the impact of antibiotics on the bacteria-phage system with multiple phage strategies and antibiotic resistance. We address conditions under which the bacteria-phage-antibiotic ecosystem results in control of the bacterial infection [14].

## PREVIOUS WORK

Many mathematical models of bacteria-phage systems exist at varying levels of complexity. The simplest models include only one phage strategy (lysis); in this simple scenario, either all bacteria are affected by the phage [39] or some bacteria are resistant to infection [40]. More complex models study the competition between two different phage strategies, such as lysis and lysogeny [41] or lysis and productive chronic infection [42]. The scope of many studies is extended to also include interactions among bacteria, phages, the host’s immune response and/or antibiotic treatment. The immune response and antibiotic agent have been modeled implicitly by modifying the rates of change of bacteria and phages [40] or explicitly by adding compartments governing antibiotic and immune response rate of change [43–45].

Other distinctions among models of bacterial infections can be made based on how bacteria reproduce. Mechanistic models incorporate a limited nutrient as an additional compartment [45–47], while more phenomenological models assume bacteria grow logistically [39, 41, 48, 49]. Furthermore, many models are used to study bacterial evolution of resistance to either phages [45, 47] or antibiotics [50]. These models are either deterministic [47] or stochastic [45, 50].

Phage and antibiotic synergy has been investigated experimentally using phage isolated from wastewater or other sources. Attention has primarily been paid to the breadth of killing that lytic phage exhibit on a diversity of *P. aeruginosa* strains, while little attention has been given to other parts of the phage lifestyle. Accordingly, models for phage-antibiotic synergy incorporate only the killing aspects of viruses [14]. These models suggest that pretreatment with phage decreases the bacteria to a low enough level that antibiotics can extinguish bacterial populations; they do not yet consider potential for phage to spread within a population and be induced by antibiotic treatment at a later time.

Consideration has been given to the impact of antibiotic treatment on the mobilization of temperate phage genetic material (including antibiotic resistance genes) between cells via transduction [51, 52]. However, to our knowledge, no mathematical models of bacteria-phage interaction have analyzed the competition between temperate and chronic phage strategies in an environment with pulses of antibiotic stress, as would happen during treatment. Filling this knowledge gap is critical to understanding the impact of antibiotic treatment on a patient infected with the bacterium *aeruginosa*.

## MODEL

### Modeling Framework

Consider a system of two competing types of bacteriophage (e.g., [41, 42]): one temperate phage *V*_*T*_ with lytic and latent lytic stages, and one chronic phage *V*_*C*_ with productive and latent stages. During the productive phase of the chronic lifestyle, phage particles are released through budding and do not kill the host bacterium. Each phage attacks one strain of bacteria that is initially susceptible to infection by either phage type. Refer to Fig 1 for an overview of the process; Fig 2 shows the complete modeling framework.

**FIG. 1.**
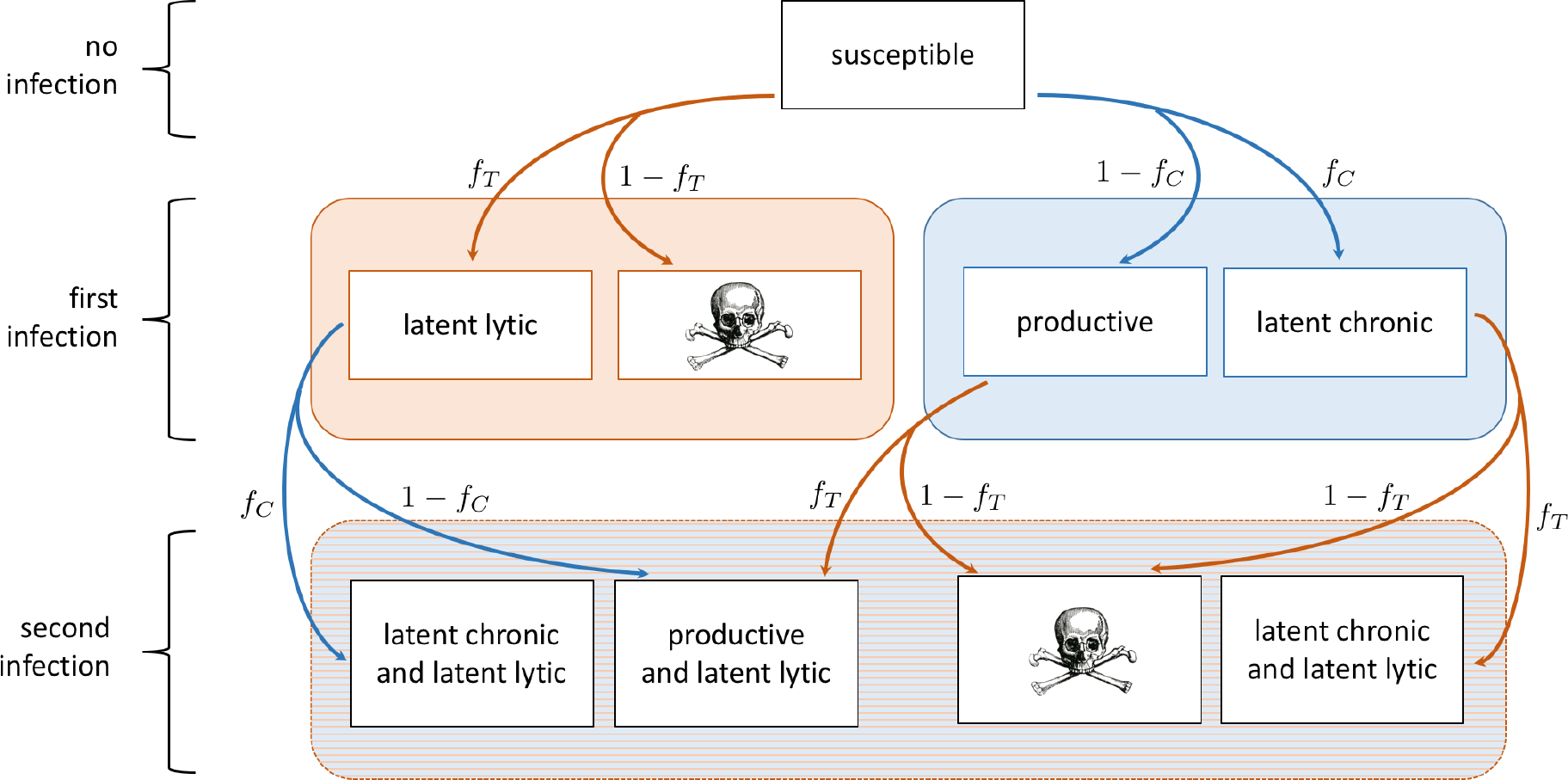
Flowchart of bacteria-phage system with both temperate (orange) and chronic (blue) phages. Boxes indicate a bacterial state, and arrows indicate an infection by phage. If a bacterium is infected by temperate phage, the probability of going latent lytic is *f*_*T*_. If a bacterium is infected by chronic phage, the probability of becoming latent chronic is *f*_*C*_. Skull sketch courtesy of Dawn Hudson (CC0).

**FIG. 2.**
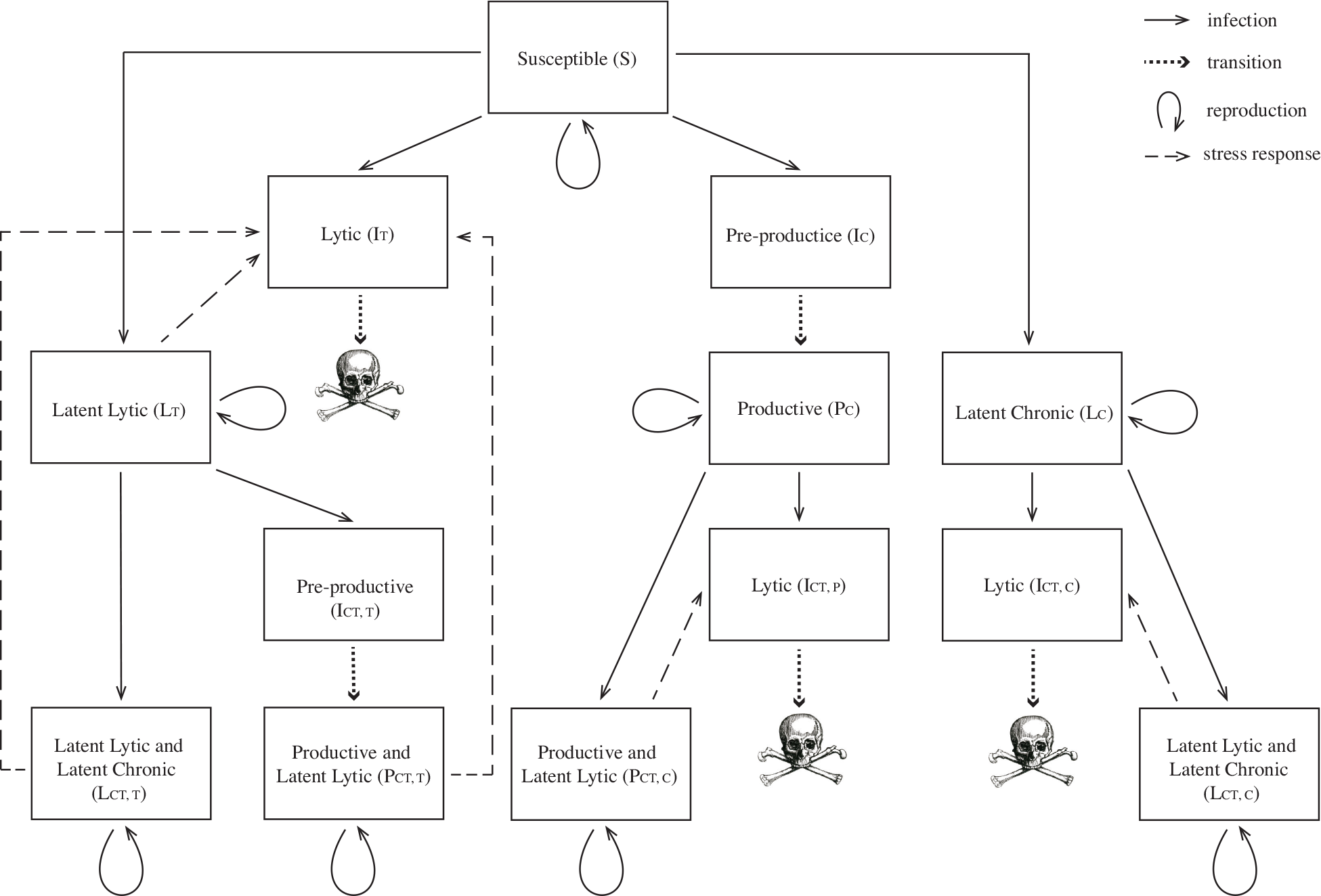
Full flowchart of bacteria-phage system, corresponding to model system (S1)–(S15). Skull sketch courtesy of Dawn Hudson (CC0).

We assume the total bacterial population *B*_tot_ grows logistically at a rate *γ* to a carrying capacity *K* [53]. Each phage infects susceptible bacteria *S* at a rate *η*. Bacteria infected by the temperate phage *V*_*T*_ will either become latently infected *L*_*T*_ with probability *f*_*T*_, or will enter a lytic state *I*_*T*_ with probability (1 − *f*_*T*_). Bacteria in the lytic state produce phage and burst (with burst size *β*_*T*_) at a rate *δ*^1^. While in the lytic state, the phage hijacks cell functions, and the cell cannot reproduce [54, 55]. Bacteria do not move between lytic and latent states unless there is a perturbation or stress to the system where viruses are induced.

Bacteria infected by the chronic phage *V*_*C*_ will either become latently infected *L*_*C*_ with probability *f*_*C*_, or will enter a pre-productive state *I*_*C*_ with probability (1 − *f*_*C*_). Bacteria in the pre-productive state stop reproducing and prepare to manufacture phage with delay rate *δ*. After the production delay, the pre-productive bacteria enter the productive state *P*_*C*_, continue reproducing at a potentially reduced rate *λ*_*γ*_, and begin producing phage at a rate *β*_*C*_ without cell death [56]. As above, after chronic phage enter the latent or productive state in a cell they will not change state. Latent chronic phage cannot be induced by stress to become productive; however, productively infected strains produce more phage under stress and reproduce more slowly. We note that biologically, productively infected strains can revert to latent infection and latent hosts can induce chronic virus production.

Once a bacterium is infected, we assume it will exclude superinfection by the same phages but may be infected by phages of the other type [57]. If a bacterium that is latently infected by the temperate phage is additionally infected with the chronic phage, the bacterium will either become latently infected with both phages 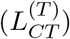 with probability *f*_*C*_, or will enter a pre-productive state 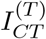 with probability (1 − *f*_*C*_). Bacteria in the pre-productive state stop reproducing and prepare to manufacture phage with delay rate *δ*. After the production delay, the infected bacteria enter the productive state 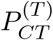, continue reproducing at a potentially reduced rate *λ*_*γ*_, and begin producing phage at a rate *β*_*C*_ without cell death [56].

Similarly, if a bacterium that is latently infected with a chronic phage is infected with the temperate phage it will either become latently infected 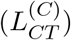 with probability *f*_*T*_, or will enter a lytic state 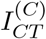 with probability (1 − *f*_*T*_). Bacteria in the lytic state produce phage and burst (with burst size *β*_*T*_) at a rate *δ*. While in the lytic state, the phage hijacks cell functions, and the cell cannot reproduce.

If a productive bacterium is then infected with the temperate phage, the bacterium will become latently infected with temperate phage 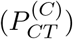 with probability *f*_*T*_. Otherwise, the productive bacterium will enter a lytic state 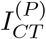 with probability (1 − *f*_*T*_). Bacteria in the lytic state produce phage and burst (with burst size *β*_*T*_) at a rate *δ*. While in the lytic state, the phage hijacks cell functions, and the cell cannot reproduce.

As shown in Fig 1, without the addition of new susceptible bacteria this infection process results quickly in a population of cells that are phenotypically either doubly infected by both phage in the latent state or are producing the chronic virus and latently infected with temperate phage.

### Infection

Many models of bacteria-phage interaction assume that a mass action process governs infection [41, 58], but *P. aeruginosa*-phage infection rates are not well-approximated by a mass action process [59, 60]. More realistically, infection rates slow as population growth activates quorum-sensing and biofilm formation [61]. One way to accommodate this infection process is to replace a mass action term with a Michaelis-Menten or Holling’s Type II functional response. In this case, all infection and absorption rates are proportional to the nonlinear response

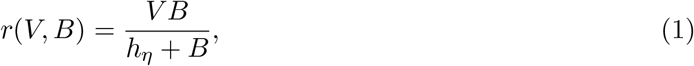

where *V* is the phage of interest, *B* is the bacteria of interest, and *h*_*η*_ is the bacteria population at which the infection rate is half of the maximum. For small bacterial populations (*B* ≈ 0), infection is approximately a mass action process. As the bacterial population grows, the infection rate saturates. See Fig 3(a).

**FIG. 3.**
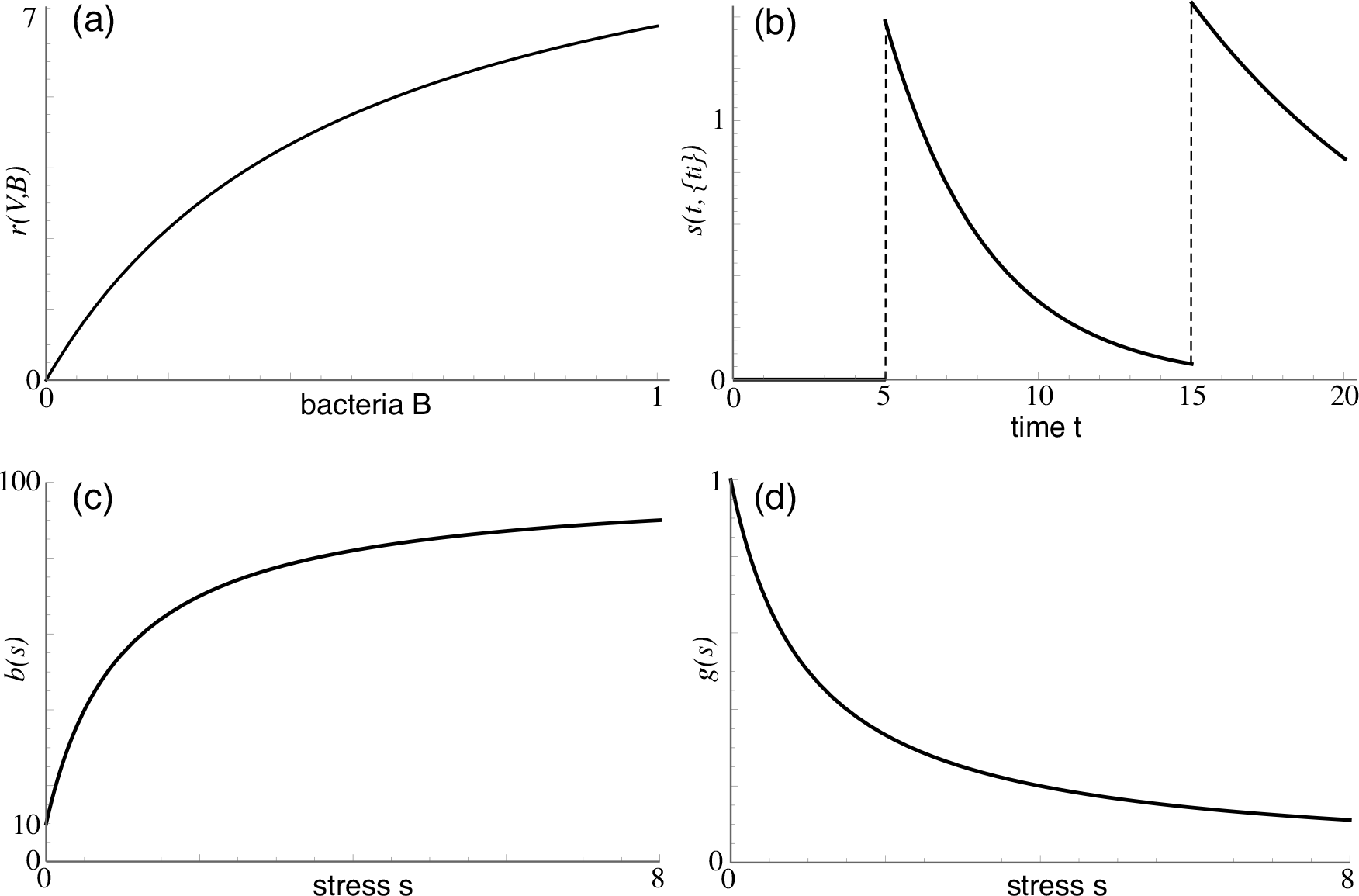
Sketches of the functions for (a) infection *r*(*V*, *B*) with phage density *V* = 10, (b) antibiotic stress *s*(*t*, {*t*_*i*_}) with {*t*_*i*_} = {5, 15}, (c) phage production *b*(*s*), and (d) cell reproduction multiplier *g*(*s*). Parameter values are taken from the baselines in Table II.

**Table I.**
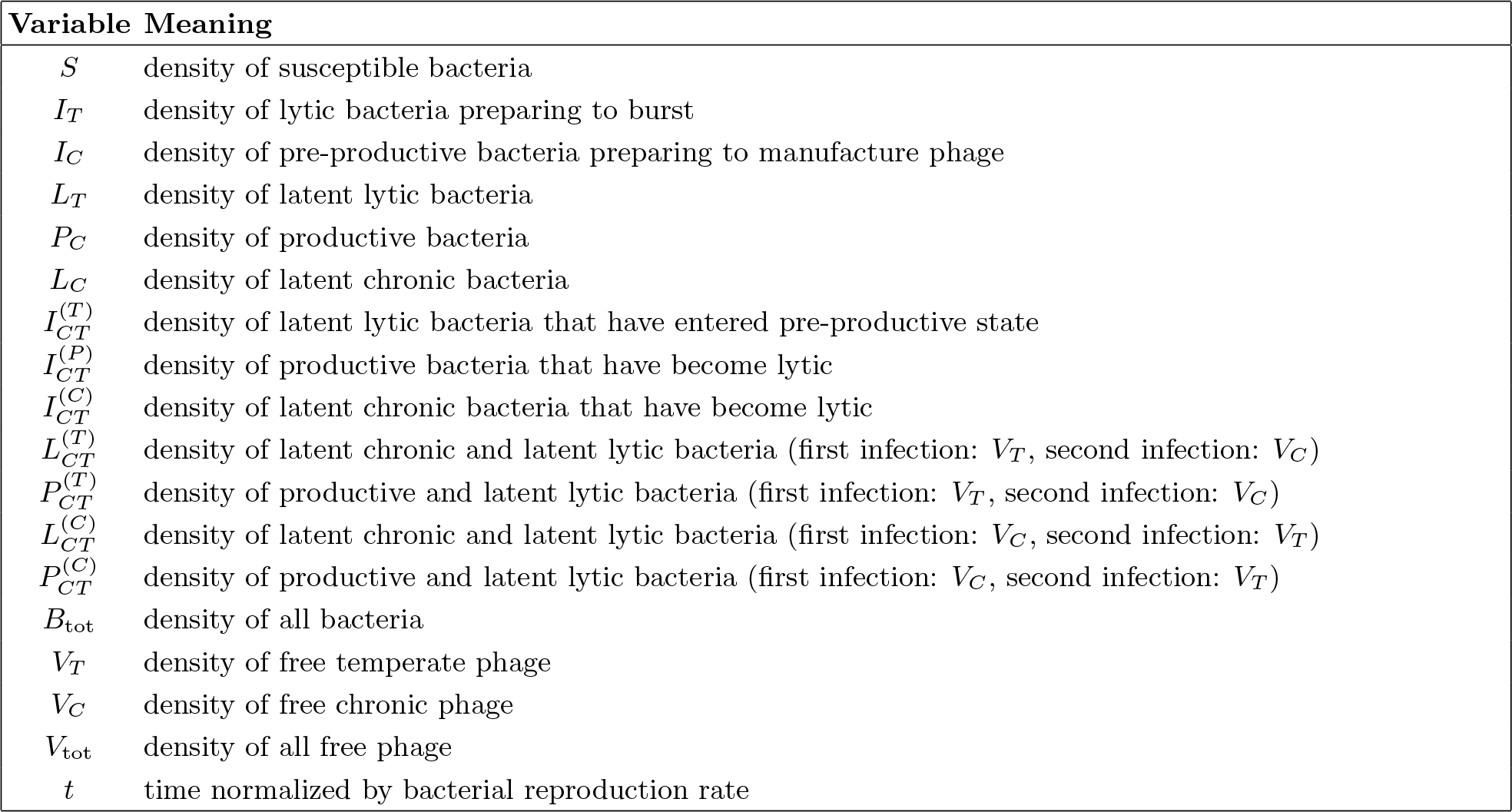
Description of model variables in bacteria-phage system (S1)–(S15). Due to nondimensionalization of density and time, all variables and parameters are nondimensional; all densities are relative to the bacterial carrying capacity and all rates are relative to the growth rate of bacteria under ideal conditions.

**Table II.** Description of model parameters in bacteria-phage system (S1)–(S15). Due to nondimensionalization of density and time, all variables and parameters are nondimensional; all densities are relative to the bacterial carrying capacity and all rates are relative to the growth rate of bacteria under ideal conditions. Commonly used time units are noted in parentheses for baseline rates.

| Parameter | Meaning | Range | Baseline | Source |
| --- | --- | --- | --- | --- |
| $\gamma$ | growth rate of bacteria under ideal conditions, normalized to 1<sup>a</sup> | 1 | 1 (5.1e-3 min<sup>-1</sup>) | [80, 81] |
| $\lambda$ | proportion growth rate change due to productive chronic infection | [0.5, 3]<sup>b</sup> | 1 | [56] |
| $K$ | carrying capacity of bacteria, normalized to 1<sup>c</sup> | 1 | 1 | [82, 83] |
| $\eta$ | infection rate | [0, 40] | 20 (0.10 min<sup>-1</sup>)<sup>d</sup> | [41] |
| $\kappa$ | bacteria death rate due to antibiotic, relative to antibiotic lysis induction rate | [0, 3.5]<sup>e</sup> | 1 | [80, 84] |
| $A$ | amplitude of stress (rate at which antibiotic induces lysis) introduced with one antibiotic dose | [0, 2] | 1.1 (5.6e-3 min<sup>-1</sup>)<sup>f</sup> | [80, 85] |
| $k$ | metabolic decay rate of antibiotic within the system | [1e-3, 0.6]<sup>g</sup> | 0.3 (1.5e-3 min<sup>-1</sup>)<sup>h</sup> | [80, 86, 87] |
| $\{t_i\}$ | vector of antibiotic administration times | — | — | [88] |
| $\delta$ | rate at which infection leads to phage production (eclipse and rise phase) | [1.5, 7.8]<sup>i</sup> | 4 (2.0e-2 min<sup>-1</sup>) | [89, 90] |
| $f_T$ | fraction of bacteria infected with $V_T$ that become latently infected | [0, 1] | 0.01 | [91, 92] |
| $f_C$ | fraction of bacteria infected with $V_C$ that become latently infected | [0, 1] | 0.01<sup>j</sup> | |
| $\beta_T$ | burst size for bacteria infected with $V_T$ | [10, 1000] | 100 | [89, 90, 93–97] |
| $\beta_C$ | phage production rate for bacteria infected with $V_C$ | [5, 200] | 10<sup>k</sup> | |
| $\beta_{\max}$ | maximum phage production rate for bacteria infected with $V_C$ under maximum stress | [10, 10000] | 100 | [34] |
| $d$ | rate of free phage degradation | [0.9, 3.6]<sup>l</sup> | 1 (5.1e-3 min<sup>-1</sup>) | [98] |

<sup>a</sup>growth rate is approximately 5.1e-3 min<sup>-1</sup> for *P. aeruginosa* grown *in vitro*, but is highly variable in cystic fibrosis patients.
<sup>b</sup>estimates based on *E. Coli* and M13 phage.
<sup>c</sup>stable bacterial density in sputum is highly variable in patients with cystic fibrosis; a study of viable *P. aeruginosa* densities in sputum of 12 patients not undergoing treatment ranged from 5.3e3 CFU/mL to 1.8e11 CFU/mL, while another study found densities between 4.02e9 CFU/g to 2.39e11 CFU/g. Log difference between control/placebo and treatment are more commonly reported.
<sup>d</sup>estimate based on *E. Coli* and $\lambda$ phage.
<sup>e</sup>estimate for antibiotic Levofloxacin (upper limit on death rate may include death by phage induction).
<sup>f</sup>estimated from *in vitro* experiment using antimicrobial peptides and meropenem.
<sup>g</sup>low estimate is for Meropenem *in vitro*, high estimate is for Ciprofloxacin *in vivo* (human).
<sup>h</sup>antibiotic is Levofloxacin (half-life approximately 6.9 hours).
<sup>i</sup>low estimate is for PAXYB1 phage and PAO1 host, high estimate is for PAK\_P3 phage and PAO1 host.
<sup>j</sup>guess based on temperate phage.
<sup>k</sup>guess based on author experience.
<sup>l</sup>low estimate is for phage extracted from Raunefjorden, high estimate is for phage extracted from Bergen Harbor (strains unknown).

### Antibiotics

Because patients infected with *P. aeruginosa* are typically treated with antibiotics at the time of bacterial detection [62, 63], we must incorporate the effects of antibiotic doses administered at times *t*_*i*_ on the bacteria-phage ecosystem. We assume that system stress spikes at times *t*_*i*_ (when antibiotics become bioavailable) and decays exponentially, consistent with typical antibiotic metabolism in the human system [64–66]. The functional form of stress is then

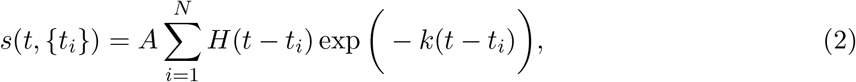

where *t* is the current time, {*t*_*i*_} is a list of antibiotic dose times, *A* is the amplitude of stress due to one antibiotic dose, *N* is the total number of antibiotic doses, *H* is the Heaviside function, and *k* is the decay rate of antibiotics in the system. For inhaled or intravenous antibiotics, the dose times are the exact times of antibiotic administration. For oral antibiotics, {*t*_*i*_} are the times at which the antibiotics become bioavailable in the blood stream. See Fig 3(b).

When the system is stressed, the following three processes occur:

1. Bacteria that are susceptible to the antibiotics die at a rate proportional to the amount of antibiotic in the system [67]. If certain strains of bacteria are resistant to antibiotics, then they will not be killed directly by antibiotics [68–70].
2. Bacteria that are infected by temperate phage induce phage production at a rate equal to the stress [34, 71, 72]. In other words, stress measures the rate at which latent lytic bacteria induce phage. Note that not all antibiotics induce phage [34], so we focus only on the types of antibiotics known to do so (e.g., quinolones like Levofloxacin and Ciprofloxacin) [8, 73]. We assume that even antibiotic-resistant bacteria induce viruses in the presence of antibiotics, which has been demonstrated for several classes of antibiotics [34, 74–77].
3. Productive bacteria increase phage production and decrease cell reproduction [78, 79]. A simple way to incorporate increased phage production during system stress is with a Hollings-like functional response. With no system stress, the phage production rate is *β*_*C*_, and with increasing system stress, the phage production rate saturates at *β*_max_:

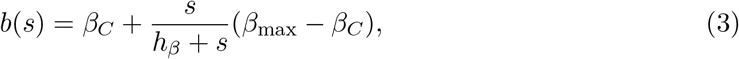

where *s* is the time-dependent stress level (2) in the system, *h*_*β*_ is the stress level at which the production rate is halfway between the minimum and maximum, and *β*_max_ is the maximum production rate when stress is maximal. See Fig 3(c). We assume that bacteria that are latently infected with the chronic virus do not induce phage production, although there is evidence that this occurs in real-world systems.

Similarly, a simple way to incorporate decreased cell reproduction during system stress is with a Hollings-like functional response. With no system stress, the cell reproduction rate is *λ*_*γ*_, and with increasing system stress, reproduction slows by a factor of *g*(*s*):
the cell eventually stops reproducing:

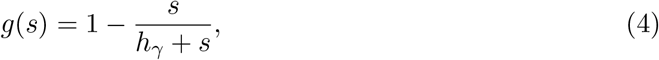

where *s* is the time-dependent stress level (2) in the system and *h*_*γ*_ is the stress level at which the growth rate is half the maximum. As stress increases, the bacterium eventually stops reproducing. See Fig 3(d).

See Tables I and II for variable and parameter definitions, respectively. See equations (S1)–(S15) for the dynamical systems model.

## RESULTS

First, we examine the model without antibiotic administration. Without external stress, the bacterial population eventually stabilizes at carrying capacity, with doubly-infected productive bacteria dominating the population (see Fig 4). Because we have assumed infection by one phage type does not prevent infection by a different type (i.e., no cross-infection exclusion) and that coinfection does not impose a fitness cost on bacteria, eventually all bacteria are infected with both phage.

**FIG. 4.**
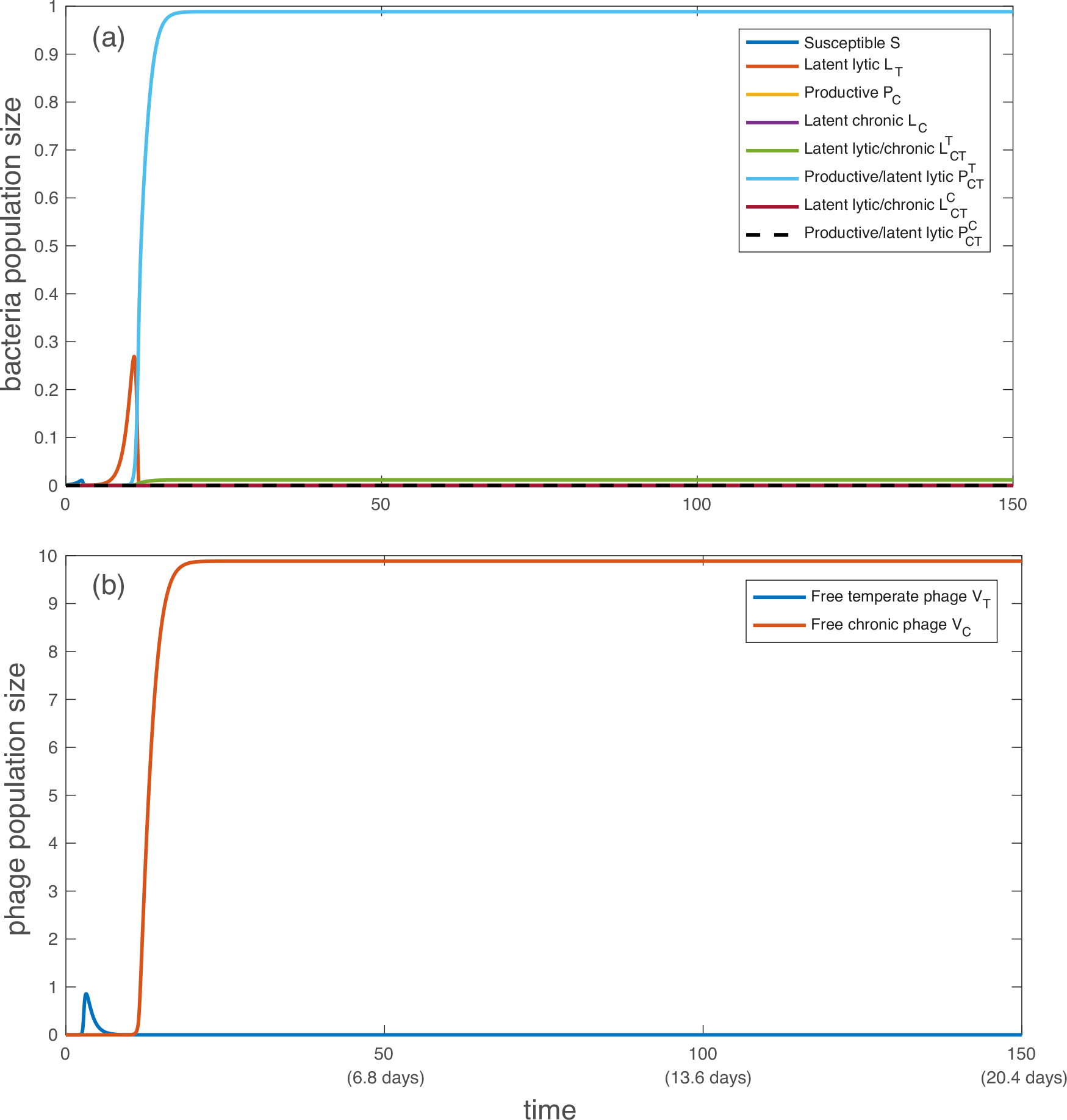
Simulation of population dynamics with no antibiotic administration: (a) bacterial population and (b) free phage population. Without antibiotics, the dominant bacterial strain is producing chronic virus while also latently infected with temperate phage 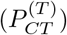, and the only free phage are chronic (*V*_*C*_). All bacteria and phage types are described in Table I. All parameter values are taken from the baselines in Table II, with *h*_*η*_ = 1/2, *h*_*β*_ = 1, *h*_*γ*_ = 1. Note that both axes are linear, not logarithmic. Initially, *S*(0) = 1e−3, *V*_*T*_ (0) = *V*_*C*_ (0) = 1e−7, following Sinha et al. [41].

Productive bacteria dominate the population because, initially, populations of bacteria latently infected with temperate phage increase faster than those latently infected with chronic phage due to the early rapid proliferation of temperate phage. Subsequently the productive strains dominate since they are formed at a much higher frequency on secondary infection than either latent infection. With a substantial population of chronically infected bacteria producing phage at steady state, the ratio of free chronic phage to bacteria stabilizes at approximately 10:1. Although little is known about the proportion of phage types seen in either clinical or wild settings, it is known that both temperate and chronic strains are often found in the same environment [99]. See Fig 5 for a visualization of the dominant path through the model system without antibiotics.

**FIG. 5.**
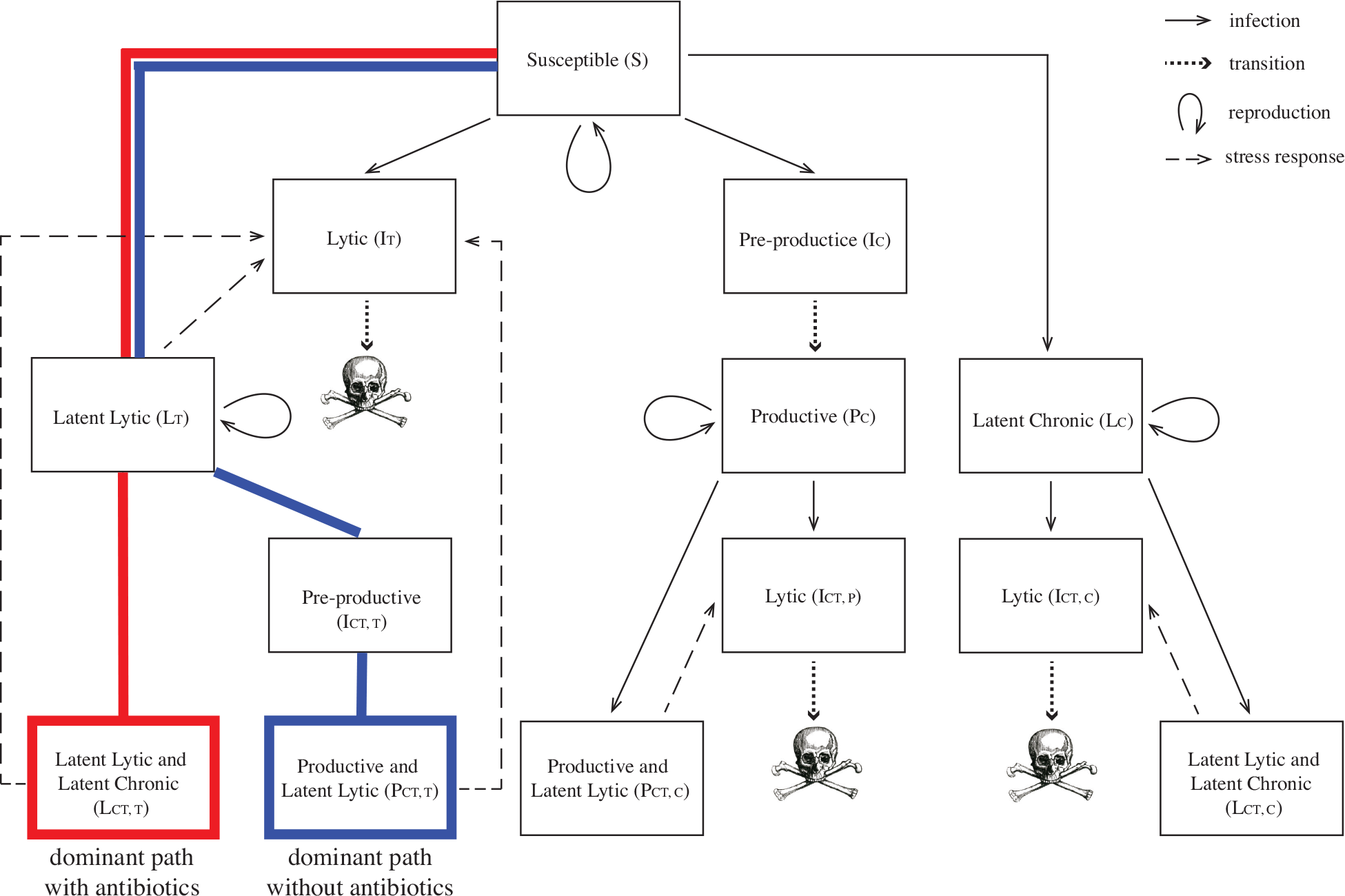
Full flowchart of bacteria-phage system, corresponding to model system (S1)–(S15), with results superimposed. The dominant path through the model compartments without antibiotics is shown in blue, while the dominant path with periodic antibiotic dosing is shown in red. Skull sketch courtesy of Dawn Hudson (CC0).

### Antibiotic treatment

Next, we examine the model where all bacteria are sensitive to antibiotics (i.e., bacteria are not resistant to the antibiotic’s intended killing mechanism, namely inhibiting bacterial DNA replication [100]) using baseline parameter values from Table II. For the purpose of illustration, we choose the period of antibiotic treatment *T* = 7.3, which is one antibiotic dose every 24 hours; this is a typical clinical dosing protocol [88]. When all bacteria are sensitive to antibiotics, periodic administration of antibiotic leads to periodic dips in bacterial populations and periodic spikes in induced free phage (see Fig 6). During antibiotic treatment, the total bacteria population remains well below the carrying capacity, and the ratio of free phage to bacteria is around 20:1 on average and about 30:1 at most. These values are consistent with existing studies of bacteria to phage ratios [101, 102].

**FIG. 6.**
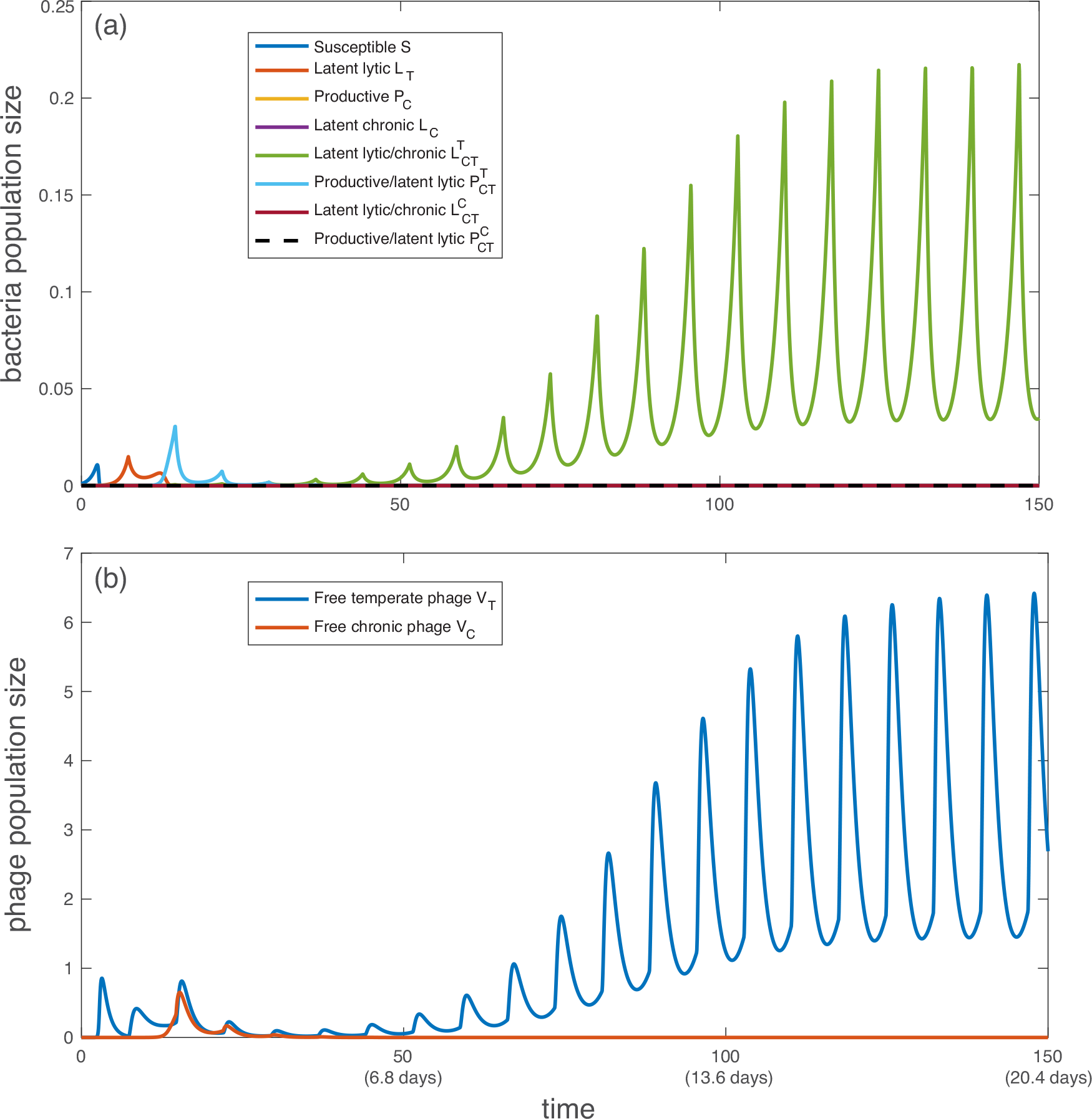
Simulation of population dynamics with no antibiotic resistance: (a) bacterial population and (b) free phage population. All bacteria and phage types are described in Table I. All parameter values are taken from the baselines in Table II, with *h*_*η*_ = 1/2, *h*_*β*_ = 1, *h*_*γ*_ = 1. Antibiotics are administered periodically every *T* = 7.3 bacterial reproductive cycles (once-daily dose). Note that both axes are linear, not logarithmic. Initially, *S*(0) = 1e−3, *V*_*T*_ (0) = *V*_*C*_ (0) = 1e−7, following Sinha et al. [41].

Fig 4 shows that without antibiotic administration, productive bacteria that are latently carrying the temperate phage are the dominant bacterial strain due to their high frequency of formation in early stages. With each antibiotic dose, the productive bacteria are replaced with strains doubly infected by latent phage, which eventually dominate the system (Fig 6). This phenomenon occurs because most bacteria that are latently infected with temperate virus (including 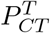) respond to antibiotic stress by inducing lysis, which brings the number of bacteria to a very low number. The drop in bacterial population allows the doubly latent infected bacteria (unencumbered by phage production) to grow slightly faster than productive bacteria and eventually dominate the population. Antibiotic administration resets the population structure from one set by initial relative frequencies of latent and active infection to one that is set by relative fitness (growth rate). The number of free chronic phage decreases over time because latently infected strains cannot become productive in this model.

To control an infection, there are two primary parameters that can be independently varied: antibiotic administration period *T* and antibiotic efficacy *κ*. The antibiotic dosing period and deadliness required to control an infection depend on other model parameters, especially the amplitude of stress caused by antibiotics and the metabolic decay rate of the antibiotic (Fig 7). Antibiotics must be administered more frequently if antibiotics are less effective at killing bacteria either directly or via induced lysis, or if antibiotics are metabolized more quickly (Fig 7a). On the other hand, antibiotics must be more effective in order to control an infection if antibiotics are administered less frequently, if antibiotic stress induces lysis less effectively, or if antibiotics are metabolized more quickly (Fig 7b). See the Supplementary Information for technical details on the sensitivity analysis.

**FIG. 7.**
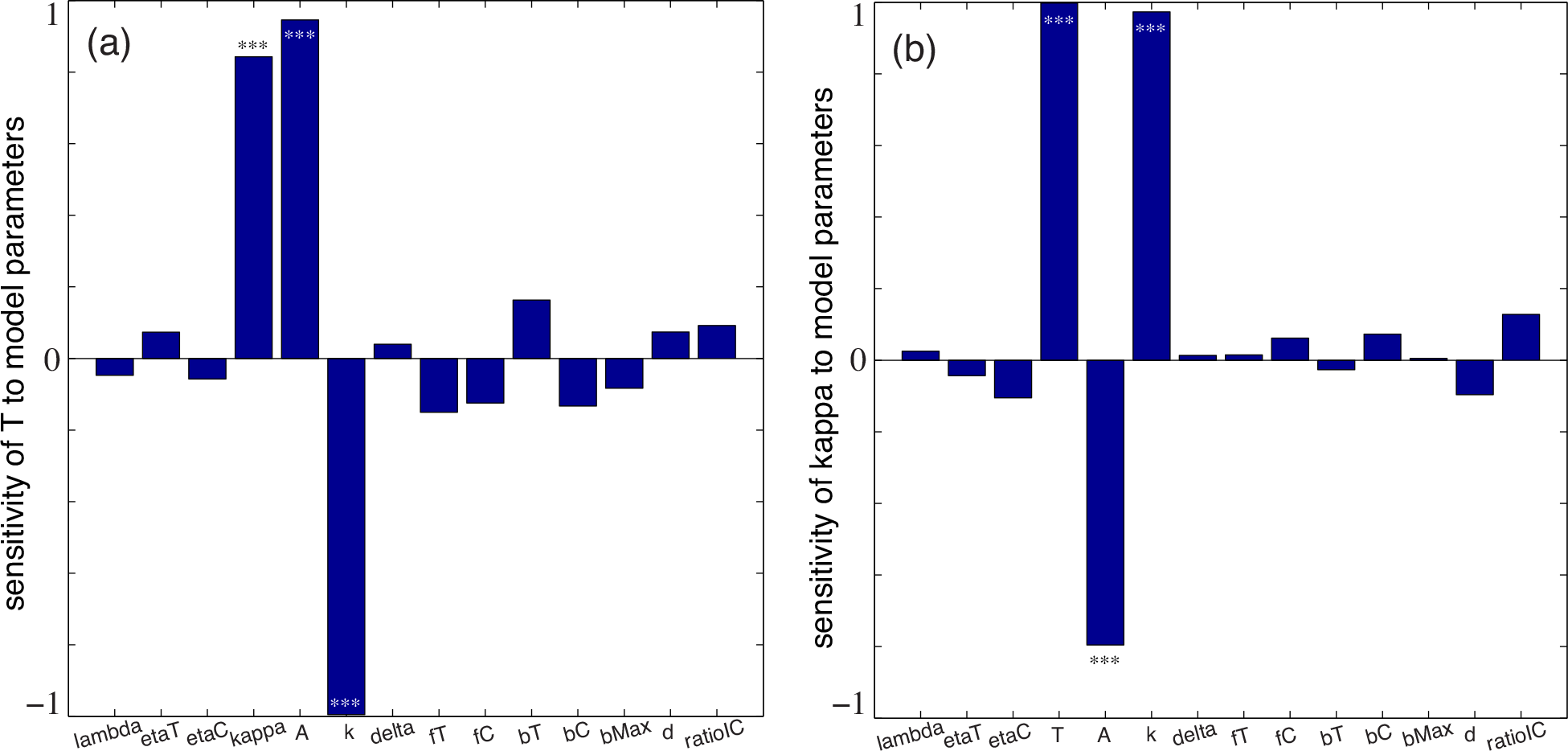
Sensitivity of (a) the antibiotic dosing period *T* required to control the infection, and (b) the antibiotic deadliness *κ* required to control the infection. The sensitivity analyses use Latin Hypercube Sampling (LHS) of parameter space and Partial Rank Correlation Coefficients (PRCC) [103]. Infection control is an average total bacterial population below 10% of carrying capacity over 300 bacterial reproductive cycles. All parameter values are taken near the baselines in Table II, with *h*_*η*_ = 1/2, *h*_*β*_ = 1, *h*_*γ*_ = 1. Initially, *S*(0) = 1e−3, *V*_*T*_ (0) = 1e−7, *V*_*C*_ (0) = ratioIC × 1e−7. The number of simulations is *n* = 150. Asterisks indicate significance (\*\*\**p* < 0.001, no asterisks *p* > 0.05). See Supplementary Information for technical details.

### Antibiotic Resistance

If all bacteria are resistant to antibiotics (*κ* = 0), then the population dynamics are qualitatively similar to when bacteria are sensitive to antibiotics. In both cases, antibiotic administration causes doubly latently infected bacteria to dominate the system. However, when all bacteria are antibiotic resistant, the total bacterial population and phage populations are noticeably larger. See Fig 8.

**FIG. 8.**
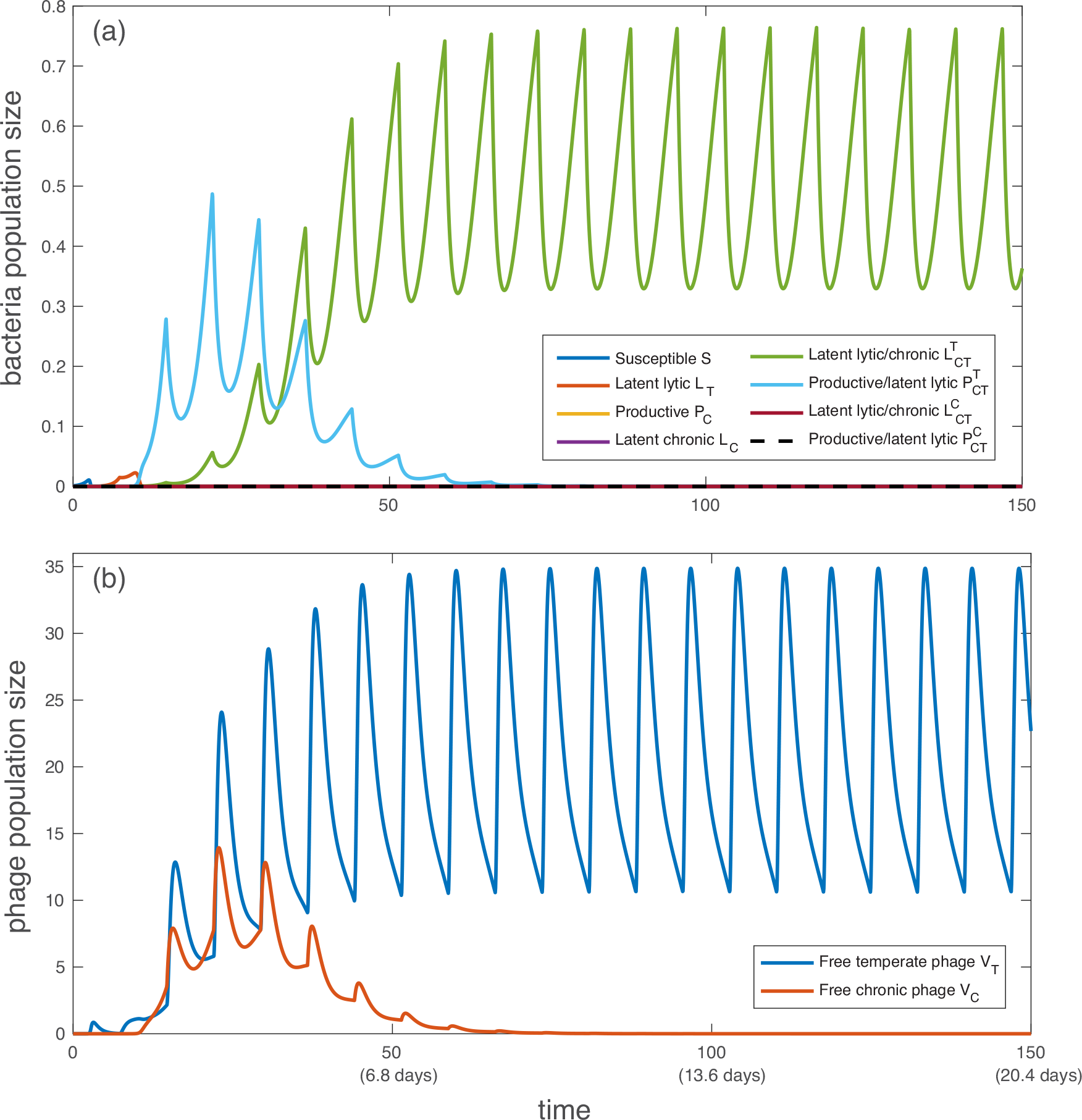
Simulation of population dynamics with complete antibiotic resistance: (a) bacterial population and free phage population. All bacteria and phage types are described in Table I. All parameter values are taken from the baselines in Table II, with *h*_*η*_ = 1/2, *h*_*β*_ = 1, *h*_*γ*_ = 1, and *κ* = 0 for all bacteria. Antibiotics are administered periodically every *T* = 10 bacterial reproductive cycles. Note that 100 bacterial reproductive cycles is approximately two weeks. Initially, *S*(0) = 1*e* − 3, *V*_*T*_ (0) = *V_C_* (0) = 1*e* − 7, following Sinha et al. [41].

### Pharmacological Implications with Antibiotic Resistance

The main concern when treating an infection with antibiotics is the size of the bacterial population. Therefore, we investigate the total bacterial population under a range of antibiotic dosing frequencies (see Fig 9). We compute the average total bacterial population over the first 300 bacterial reproductive cycles (40.8 days), and we find that both antibiotics and temperate phage are critical to controlling the infection and work synergistically even when bacteria are antibiotic-resistant. We define infection control to be an average bacterial population less than 10% of carrying capacity (i.e., 1-log decrease in bacteria levels compared with placebo).

**FIG. 9.**
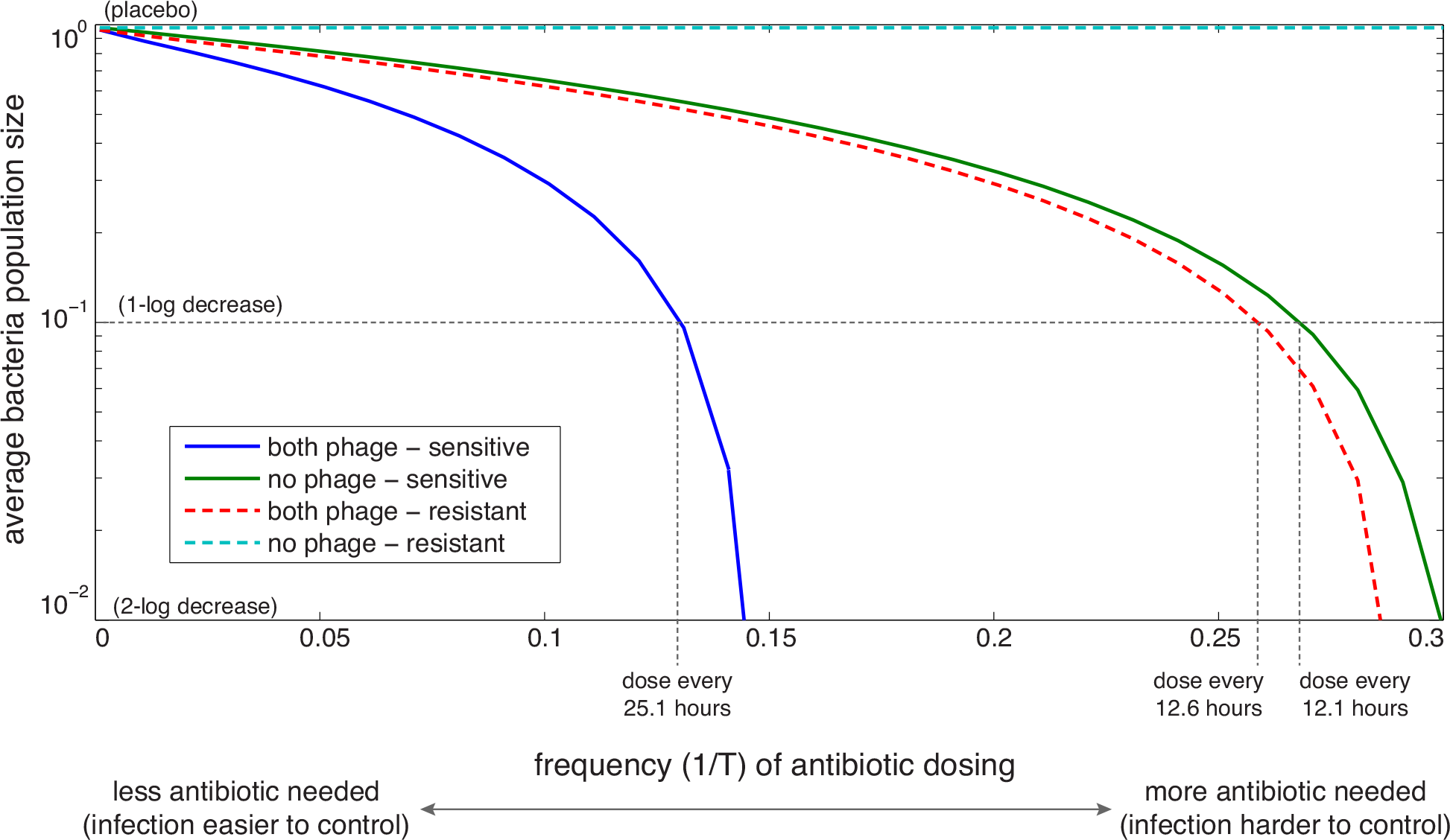
Average total bacterial population for a range of periodic antibiotic dosing protocols. All parameter values are taken at the baselines in Table II, with *h*_*η*_ = 1/2, *h*_*β*_ = 1, *h*_*γ*_ = 1, *t*_max_ = 300. Solid lines indicate that all bacteria are sensitive to antibiotics, and dashed lines indicate that all bacteria are resistant. Note that the vertical axis is logarithmic, while the horizontal axis is linear. Initially, *S*(0) = 1*e* − 3, *V*_*T*_ (0) = *V*_*C*_ (0) = 1*e* − 7, unless otherwise noted.

If only chronic phage are present in the system (Fig S1a), effective antibiotics are required to control the infection. If all bacteria are sensitive to antibiotics, the presence of chronic phage controls the infection slightly better than if there are no chronic phage due to cost of production during productive infection.

If only temperate phage are present in the system (Fig S1b), infection is controlled even when bacteria are resistant. In fact, the efficacy of temperate phage alone is similar to the efficacy of antibiotics alone. With both effective antibiotics and temperate phage, the number of antibiotic doses required to keep the infection under control is cut in half compared with antibiotics alone or temperate phage alone.

If both phages are present in the system (Fig 9), infection control is marginally better than if only temperate phage are present (Fig S1b). These results demonstrate the synergy between temperate phage and antibiotics even in resistant populations. No deliberate combination therapy may be needed to treat these infections because temperate phage are commonly found in natural populations of *P. aeruginosa* bacteria [99].

## DISCUSSION

The model presented here shows that temperate phage infection makes antibiotic treatment of bacterial infections both more effective and efficient, whether or not the bacteria are susceptible to the antibiotics. When bacteria are sensitive to antibiotics, then antibiotic treatments need not be as frequent if temperate phage are present. Even if some or all bacterial strains are antibiotic resistant, antibiotics may still be able to control the infection in the presence of phages by triggering phage induction and cell lysis. For the rest of the discussion, we will assume that an infection is controlled if the average total bacterial population remains below 10% of carrying capacity over 300 bacterial reproductive cycles; in clinical terms, control is a 1-log difference between *P. aeruginosa* density in sputum for patients given antibiotics versus placebo over 40.8 days.

For *P. aeruginosa* bacterial infections that respond to antibiotics, the model predicts that standard antibiotic doses need to be administered approximately every 12.1 hours if no phage are present but only once every 25.1 hours if temperate phage are present (see Fig 9). If bacteria are all antibiotic resistant, then temperate phages are required to control the infection, and antibiotic dosing is required every 12.6 hours to sufficiently induce lysis.

These findings are consistent with clinical evidence; patients with cystic fibrosis (CF) given aerosolized Levofloxacin twice-daily experienced a nearly 10-fold decrease in *P. aeruginosa* density (our definition of infection control) over the treatment period compared with the placebo group [104]. The study did not investigate the presence of phage, but did note that approximately 60% of *P. aeruginosa* isolates showed resistance to Levofloxacin, supporting our prediction that dosing should fall between once- and twice-daily depending on the susceptibility of the bacteria to antibiotics. Our findings are also consistent with existing antibiotic dosing protocols; although aerosolized quinolones are no longer approved for CF patients, IV and oral doses are commonly recommended on a once-, twice-, or thrice-daily schedule [88, 105].

While chronic phages are marginally beneficial in controlling infections, they are not able to control an infection without either temperate phages or effective antibiotics. In fact, chronic phages may actually inhibit control of infections by disrupting the human immune response [106, 107], a detail not yet incorporated into our model.

Like all models, our model has limitations. In the interest of simplicity, we have ignored the possibility of multiple infections by the same phage type. However, many phages that infect *P. aeruginosa* produce superinfection exclusion proteins that effectively prevent multiple infections by the same phage type [77, 108]. We also do not include the exclusion of one phage type by the other. Little is known about cross resistance to phage infection; it is often assumed to be uncommon, but including cross resistance may dramatically impact the model predictions. If cross resistance is in fact common, it is possible that phage-antibiotic synergy breaks down for some range of model parameters; we leave this analysis for future study.

Also, our model assumes that antibiotics induce phage, so this model is only applicable with quinolone antibiotics like Levofloxacin and Ciprofloxacin [34]. However, drugs from this class of antibiotics are commonly used to treat *P. aeruginosa* infections [104, 109].

In addition, some phage are able to detect bacteria population density, which appears to affect the frequency of lysogeny [110, 111]. If this process applies to *P. aeruginosa* and its phages, a more sophisticated model would incorporate a density-dependent latency probability: *f*_*T*_ (*B*_tot_) and *f*_*C*_ (*B*_tot_).

The model additionally assumes that bacteria resistant to antibiotics are still susceptible to lysis via phage induction, but this phenomenon depends of the mechanism of antibiotic resistance. There are many mechanisms of resistance to quinolones and fluoroquinolones. However, sub-inhibitory concentrations of several antibiotics are known to induce SOS but not result directly in cell death [34, 74–77]. Therefore we model the impact of phage induction on *P. aeruginosa* population size with and without antibiotic resistance.

Because this model does not include an evolutionary dynamics component, the results presented here are only applicable to acute exacerbations. If bacteria/phage evolution were integrated into this model, it may be able to explain longer-term dynamics seen in chronic infections in humans [101].

Also, all latent chronic infection states are final such that virus production cannot be induced by stress. We believe changing the model structure to accommodate chronic phage induction may change the number of productive bacteria but would not change the overall impact of antibiotic synergy, which primarily occurs with temperate infections.

Finally, the quantitative results presented in Fig 9 depend significantly on how effective antibiotics are at killing bacteria directly versus killing via phage induction (*κ* in our model). To our knowledge, no study has experimentally measured the relative number of bacteria killed by the intended antibiotic mechanism versus phage induction, so we assume that antibiotics kill via each method equally quickly. If antibiotics directly kill bacteria much more quickly (*κ* > 1), then antibiotic resistance is more detrimental to infection control than lack of phages. If antibiotics trigger phage induction much more quickly (*κ* < 1), then a lack of phages is more detrimental to infection control than antibiotic resistance. Experimental work is needed to determine a reasonable range for *κ* and test whether it is an evolvable trait.

## CONCLUSION

Antibiotic resistance threatens the efficacy of standard treatments for many dangerous and common infections. Using *P. aeruginosa* infections as motivation, we present a theoretical case for using antibiotics that trigger phage induction (e.g., quinolones) to treat bacterial infections. We show that if bacteria are antibiotic resistant, then using antibiotics in the presence of phages can still control the infection. If bacteria are susceptible to antibiotics, then the presence of phages allow for less frequent antibiotic dosing, which reduces the risk for antibiotic resistance in the future. In either case, the natural presence of phages in bacterial populations allows for more effective treatment of common bacterial infections. These, strain-dependent responses to antibiotics suggest the importance of personalized medicine approaches to treatment of infectious disease.

As a final perspective, we remember that phage induction and bacterial death may occur across the microbiome of individual hosts treated with antibiotics. The impact of these dynamics in a community context must be considered carefully on the stability of the microbiome ecosystem as a whole.

## ACKNOWLEDGEMENTS

This work was funded in part by the National Science Foundation grant DMS-1815764 (ZR), the Cystic Fibrosis Foundation grant WHITAK16PO (RJW), an Allen Distinguished Investigator Award (RJW), and National Institutes of Health grant R37 AI83256-06 (GAO). The funders had no role in study design, data collection and analysis, decision to publish, or preparation of the manuscript.

## SOFTWARE AVAILABILITY

All software (Matlab .m files) is publicly available from the Illinois Data Bank: https://databank.illinois.edu/datasets/IDB-9721455 [112].

## SUPPLEMENTARY INFORMATION

### Model

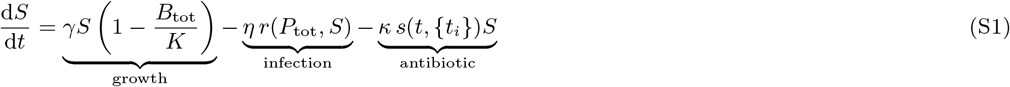

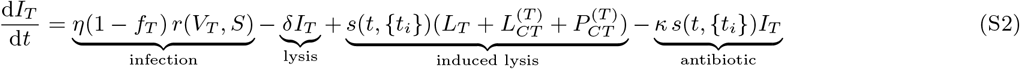

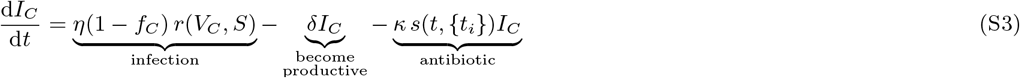

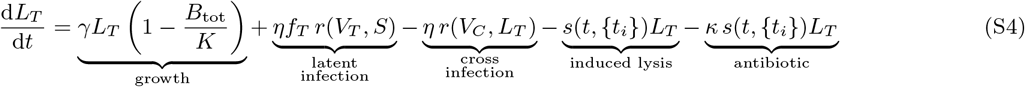

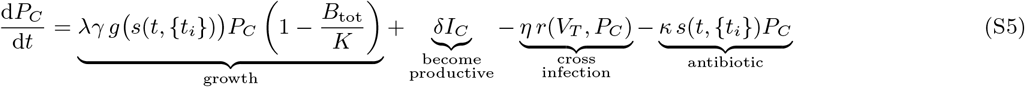

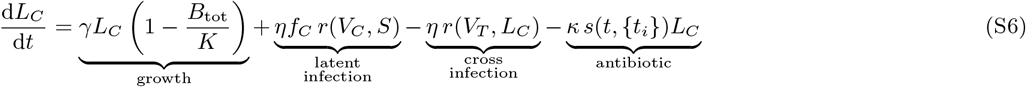

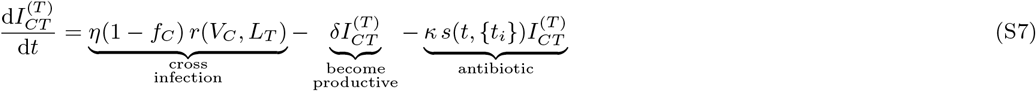

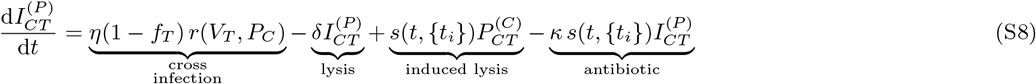

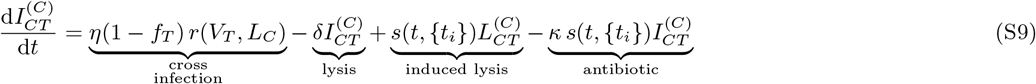

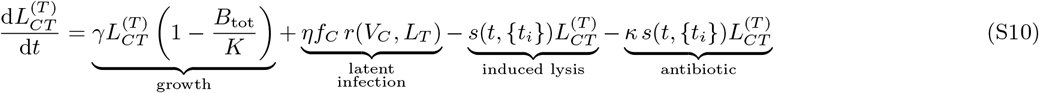

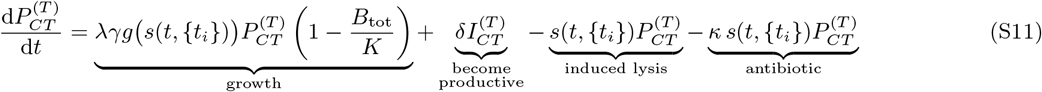

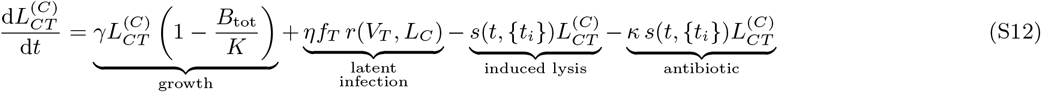

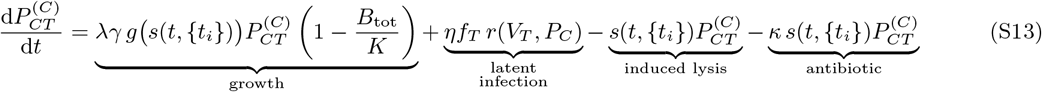

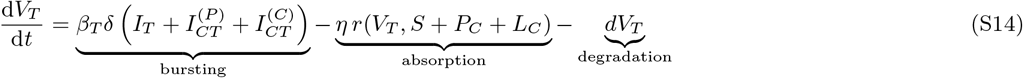

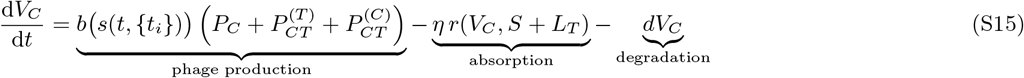

### Sensitivity analysis: technical details

In all analyses, we have used the initial conditions *S*(0) = 1e−3, *V*_*T*_ (0) = *V*_*C*_ (0) = 1e−7, following Sinha et al. [41], unless otherwise noted. While the steady state dynamics are not identical under different initial conditions, we note that the qualitative steady state behaviors (e.g., surviving strain types, dominant phage strategies, peak bacterial population sizes during antibiotic treatment, etc.) are robust to initial condition changes. Also, we have demonstrated in Fig 7 that the initial ratio of phage types does not significantly impact either the minimum antibiotic dosing period or the minimum deadliness needed to control the infection.

The global uncertainty and sensitivity analysis was performed using the methodology outlined in Marino et al. [103]. The base code for the analysis is freely available at the author Denise Kirschner’s website [113]. The specific implementation is available at the Illinois Data Bank repository: doi.org/10.13012/B2IDB-9721455_V1 [112].

In brief, the analysis uses Latin Hypercube Sampling (LHS) of parameter space to simulate uncertainty in model parameters. LHS sampling requires fewer model simulations than simple random sampling without introducing bias [114]. We used uniform sampling of each parameter about the base values given in Table II. The ranges of the uniform samples are available in our code.

We use Partial Rank Correlation Coefficients (PRCC) to test the sensitivity of model outputs to parameter uncertainty because model outputs generally depend monotonically on model inputs, but those relationships are not linear trends. As noted by Marino et al. [103], for linear trends we could have used Pearson correlation coefficient (CC), partial correlation coefficients (PCCs), or standardized regression coefficients (SRC). Had our trends been non-monotonic, we would have used the Sobol method or one of its many extensions [115].

The displayed sensitivities in Fig 7 are the PRCCs for the model output *y* (either minimum *T* or minimum *κ*) and the model inputs *x*_*j*_. As described by Marino et al. [103], partial rank correlation characterizes the monotonic relationship between input *x*_*j*_ and output *y* after the effects on *y* of the other inputs are removed. The values of PRCCs fall between −1 and 1, with 1 indicating the strongest positive rank correlation and −1 indicating the strongest negative rank correlation. The significance indicates the probability that the rank correlation is zero (i.e., large significance suggests that there is no relationship between *x*_*j*_ and *y*).

## Supplemental results

**FIG. S1.**
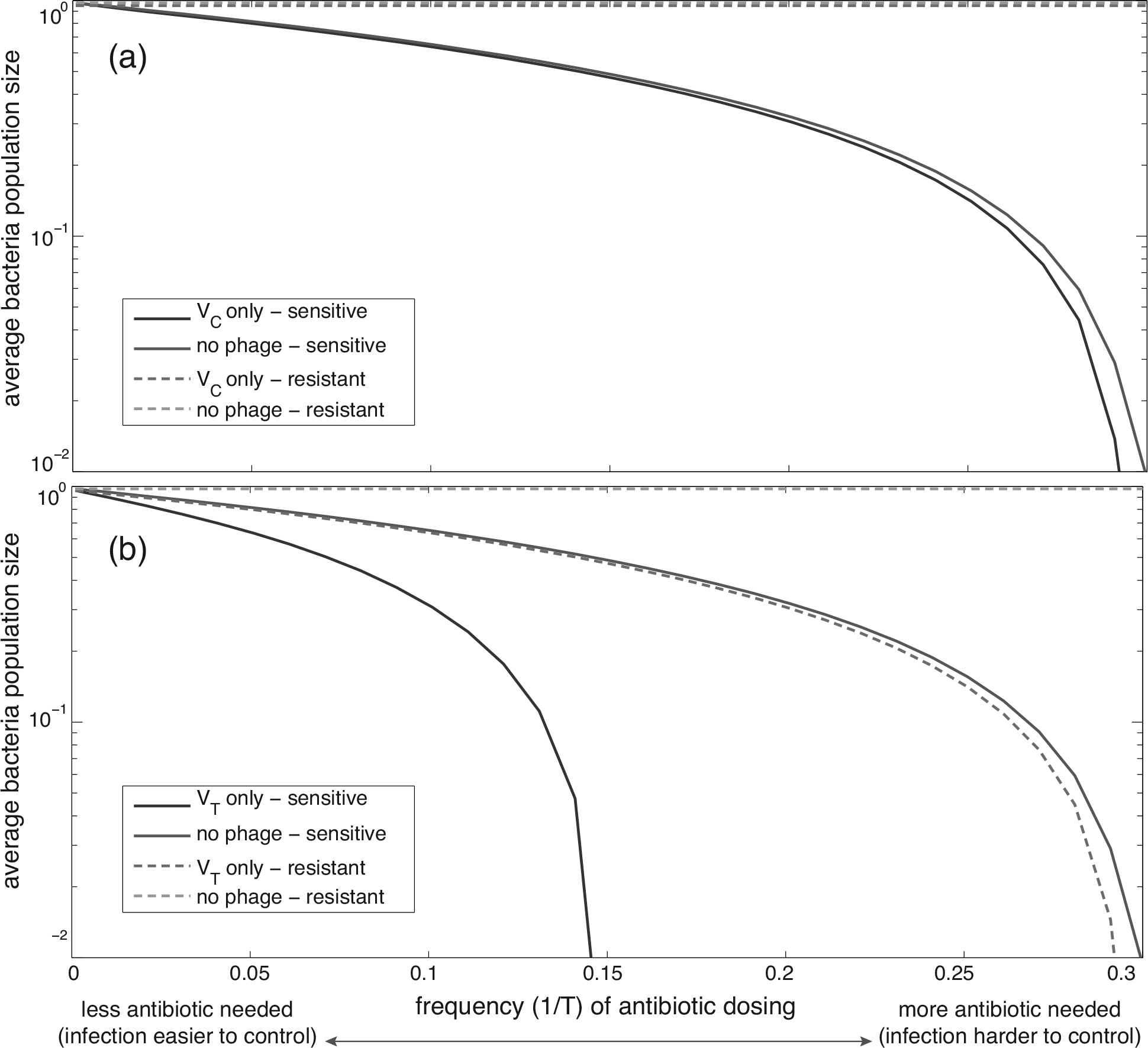
Average total bacterial population for a range of periodic antibiotic dosing protocols and only one type of phage. (a) No temperate phage present in system. (b) No chronic phage present in system. All parameter values are taken at the baselines in Table II, with *h*_*η*_ = 1/2, *h*_*β*_ = 1, *h*_*γ*_ = 1, *t*_max_ = 300. Solid lines indicate that all bacteria are sensitive to antibiotics, and dashed lines indicate that all bacteria are resistant. Initially, *S*(0) = 1e−3, *V*_*T*_ (0) = *V*_*C*_ (0) = 1e−7, unless otherwise noted.

## Footnotes

1 This modeling choice circumvents the necessity of a delay differential equation.

